# Purifying selection and adaptive evolution proximate to the zoonosis of SARS-CoV-1 and SARS-CoV-2

**DOI:** 10.1101/2023.08.07.552269

**Authors:** Jeffrey P. Townsend, Stephen Gaughran, Hayley B. Hassler, J. Nicholas Fisk, Mofeed Nagib, Yinfei Wu, Yaning Wang, Zheng Wang, Alison P. Galvani, Alex Dornburg

## Abstract

Over the past two decades the pace of spillovers from animal viruses to humans has accelerated, with COVID-19 becoming the most deadly zoonotic disease in living memory. Prior to zoonosis, it is conceivable that the virus might largely be subjected to purifying selection, requiring no additional selective changes for successful zoonotic transmission. Alternatively, selective changes occurring in the reservoir species may coincidentally preadapt the virus for human-to-human transmission, facilitating spread upon cross-species exposure. Here we quantify changes in the genomes of SARS-CoV-2 and SARS-CoV-1 proximate to zoonosis to evaluate the selection pressures acting on the viruses. Application of molecular-evolutionary and population-genetic approaches to quantify site-specific selection within both SARS-CoV genomes revealed strong purifying selection across many genes at the time of zoonosis. Even in the viral surface-protein Spike that has been fast-evolving in humans, there is little evidence of positive selection proximate to zoonosis. Nevertheless, in SARS-CoV-2, NSP12, a core protein for viral replication, exhibited a region under adaptive selection proximate to zoonosis. Furthermore, in both SARS-CoV-1 and SARS-CoV-2, regions of adaptive selection proximate to zoonosis were found in ORF7a, a putative Major Histocompatibility Complex modulatory gene. These findings suggest that these replication and immunomodulatory proteins have played a previously underappreciated role in the adaptation of SARS coronaviruses to human hosts.

## Introduction

The COVID-19 pandemic resulted from the third time in the past two decades that a coronavirus has zoonosed into humans and caused severe disease (Zhang and Holmes 2020). The SARS-CoV-1 epidemic was contained (Wilder-Smith and Freedman 2020) and the ongoing MERS-CoV viruses exhibit low transmissibility between humans (Zhang and Holmes 2020). In contrast, SARS-CoV-2 has claimed over six million lives since 2019 (World Health Organization 2023). Epidemiologists have long predicted that human population growth and consequent interactions with wildlife will increase the rate of viral zoonotic transmission (Patz et al. 2004; Wolfe et al. 2005; Wood et al. 2012; Plowright et al. 2015; Galvani et al. 2016). Molecular adaptations that facilitate zoonosis may be rare: strong purifying selection dominates genome evolution in clade-based analyses of Severe Acute Respiratory Syndrome (SARS) viruses in reservoir populations (Li et al. 2020). However, our understanding of the evolutionary dynamics of zoonotic viruses is still nascent (Donato and Vijaykrishna 2017; Olival et al. 2017; Wang et al. 2020; Valero-Rello and Sanjuán 2022). Therefore, it remains unclear whether chance wildlife interactions alone mediate zoonosis or whether adaptive evolution at the time of zoonosis is a core component of zoonotic transmission (Holmes and Drummond 2007; Holmes 2009; Engering et al. 2013; Santos et al. 2023).

Hundreds of viral genomes were sequenced during the 2002 SARS-CoV-1 epidemic. Throughout the COVID-19 pandemic, millions of SARS-CoV-2 genomes were sequenced. Evolutionary models can be applied to extensive viral sequence data to detect signals of positive and negative selection (Bhatt, Holmes & Pybus 2011, Vitti *et al*. 2013, Park *et al*. 2015). These coronavirus sequences have provided invaluable data and insights on the molecular evolution of variants that have arisen after zoonosis (Emam et al. 2021; Kumar et al. 2021; Rochman et al. 2021; Xi et al. 2022). SARS-CoV-1 and SARS-CoV-2 sequences have been leveraged to evaluate selection at deep timescales, (inclusive of both human and wildlife coronaviruses; Forni et al. 2017; MacLean et al. 2021; Morales et al. 2021). These analyses reveal strong signatures of purifying selection: numerous amino-acid residues are identical among coronaviruses circulating within reservoir bat populations (Li et al. 2020; MacLean et al. 2021). Some of these conserved sequences encode proteins that interact with the host ACE2 receptor; some feature O-linked glycans that could facilitate immune evasion (Andersen et al. 2020). Analysis of numerous insertions or disparities in sequence identity occurring subsequent to zoonosis has indicated positive selection in human hosts (Cagliani et al. 2020; van Dorp et al. 2020; Obermeyer et al. 2022). For example, Anderson et al. (2020) revealed a unique polybasic cleavage site in sequences of human SARS-CoV-2 spike proteins that potentially augments its capability to infect humans and transmit after infection. In addition, recent investigations of selection in the Omicron strain have revealed clusters of mutations that alter Spike protein functionality and enable escape from neutralizing antibodies (Martin et al. 2022). However, selective pressures encountered along the phylogenetic branch proximate to zoonosis remain unclear because of differential surveillance in reservoirs versus human hosts as well as the narrow timescale over which selection during zoonosis must be resolved.

Here we analyze selection on SARS-CoV-2 and SARS-CoV-1 viral genomes around the time of zoonosis to provide insight into the adaptive and non-adaptive processes that govern the process of zoonosis. To quantify patterns of site- and region-specific selection across the SARS-CoV-1 and SARS-CoV-2 genomes around the times of their zoonoses, we apply a model that provides power to resolve selection at a narrow molecular evolutionary time scale. Our analyses reveal the gene regions under especially strong purifying or positive selection during zoonosis, thereby providing crucial molecular evolutionary context regarding the inter-species transmissibility of coronaviruses from wildlife to humans.

## Methods

### Data acquisition and phylogenetic analyses

We acquired 559 viral genome sequences from GSAID (Elbe and Buckland-Merrett 2017; Shu and McCauley 2017), including 130 human SARS-CoV-1, 373 human SARS-CoV-2, and 56 bat/pangolin non-fragmented SARS-like sequences. Nucleotide sequences were aligned using `MAFFT` (Katoh and Standley 2013) and `Translator X` (Abascal et al. 2010) guided by the corresponding amino-acid alignment. Sequences corresponding to the *E*, *M*, *N*, ORF1a, ORF1b, ORF3a, ORF6, ORF7a, ORF7b, ORF8, and *S* genes were extracted from the viral genomic alignment. All gene sequence alignments were validated by eye to ensure automated approaches did not lead to erroneous statements of homology.

To estimate the molecular phylogeny of SARS-CoV-1, SARS-CoV-2 and SARS-like viruses from reservoir species, we used two maximum-likelihood approaches. First, we analyzed a concatenated alignment using `RAxML` v7.2.8 (Stamatakis 2014), specifying a General Time-Reversible (GTR) model of nucleotide substitution with a discretized gamma-distributed model of rate heterogeneity. Second, we used `IQ-TREE 2` conditioning on the best-fit model and the partitioning strategy; the best-fit model was identified using the Bayesian Information Criterion (Minh et al. 2020) to assess the robustness of our results to topological uncertainty across the two likelihood search algorithms, as well as model and partition choice (Kainer and Lanfear 2015). To estimate relative divergence times using the maximum-likelihood trees and virus sampling dates, we employed RelTime with Dated Tips (RTDT; Miura et al. 2020). This chronogram was used as a guide tree for the estimation of site-rates using a custom implementation of `HyPhy` (Pond et al. 2005) within `PhyDesign` (López-Giráldez and Townsend 2011). Using the resulting site rates in `PhyInformR` (Dornburg et al. 2016), profiles of phylogenetic informativeness (Townsend 2007) were generated, enabling graphical depiction of the molecular evolutionary information in each gene for the phylogenetic branches under investigation for molecular evolutionary selection.

### Ancestral genome reconstruction

To provide critical historical context for understanding the evolution of zoonosis, ancestral nucleotide sequences for each gene were inferred using the RAxML tree topology. Specifying a generalized time-reversible substitution model with a gamma distribution of rates, we reconstructed ancestral sequences of the most recent common ancestors of viruses infecting humans and the most closely related reservoir strain for both SARS-CoV-1 and SARS-CoV-2 using maximum-likelihood via `FastML` (Ashkenazy et al. 2012). To assess the robustness of our results to an alternate approach for ancestral sequence reconstructions, we repeated these analyses using the empirical Bayes ancestral state reconstruction algorithms in `IQ-TREE 2` (Minh et al. 2020).

### Identifying selection proximate to zoonosis using MASS-PRF

We used `MASS-PRF` (2017) to profile the strength of selection in the phylogenetic branches giving rise to the zoonotic progenitors of SARS-CoV-1 and SARS-CoV-2 across the genomes of SARS-CoV-1 and SARS-CoV-2. `MASS-PRF` averages clustered-site Poisson Random Field (PRF) population genetic polymorphism and divergence models of selection (Sawyer and Hartl 1992), analyzing divergent sequences between taxa and polymorphic sequences within taxa. Selection proximate to zoonosis can be assessed by comparative analysis of post-zoonotic polymorphism versus mutations that were fixed in the pre-zoonotic period immediately prior to emergence in humans. We designated all sequence variants in our dataset from SARS-CoV-1 or SARS-CoV-2 as polymorphic, and assessed divergence from the reconstructed sequence of the ancestral node between the human sequences (i.e. SARS-CoV-1 or SARS-CoV-2) and the most proximate non-human virus sequences.

We analyzed each gene separately. For the polyprotein genes ORF1a and ORF1b, we analyzed each of 16 constituent non-structural proteins (NSP) separately. To obtain profiles of selection, we averaged 30,000 randomly sampled `MASS-PRF` models. Site-specific divergence times and mutation rates (akin to phylogenetic branch lengths) were estimated based on synonymous polymorphism and divergence. 95% model intervals for the scaled selection coefficient 𝛄 were identified and plotted, illustrating gene regions and sites where the intensity of selection was significantly different from neutrality (𝛄 = 0). Criteria for selection strength were determined as in Zhao et al. (2017): positive selection was deemed strong when 𝛄 > 4 and weak when 4 > 𝛄 > 0. Statistical significance of positive selection was determined when the lower bound of the 95% model interval was greater than zero. Weak negative selection was indicated when 0 > 𝛄 > −1. Strong negative selection was indicated when 𝛄 <−1. Statistical significance of negative selection was determined when the upper bound of the 95% model interval was less than 0. Note that the incommensurate thresholds (4 and −1) arise because 𝛄 scales asymmetrically over neutrality at 𝛄 = 0. Any gene exhibiting insufficient divergence to enable quantitative estimation of 𝛄 was assumed not to have experienced positive selection, since positive selection is expected to lead to substantial molecular divergence (Zhao et al. 2017).

### Identifying post-zoonotic selection using PAML

We used `codeml` in the `PAML` package (Yang 2007) to quantify the posterior probability of positive selection following zoonosis across the genomes of SARS-CoV-1 and SARS-CoV-2. We applied the site model in `codeml` to sequences from two clades: all SARS-CoV-1 sequences and all SARS-CoV-2 sequences. For each clade, we used `IQ-TREE 2` (Minh et al. 2020) to construct maximum-likelihood phylogenetic trees for each gene and NSP using a GTR substitution model with gamma rate heterogeneity. From these phylogenies, we ran the M8 (beta and *ω*) site model in `codeml` (Yang et al. 2000), which allows *ω* to vary among sites across the gene. We applied the Bayes empirical Bayes method (Yang et al. 2005) embedded in the PAML package to estimate the posterior probabilities of sites under positive selection. Sites with posterior probability > 0.80 were reported, though we only deemed those sites with posterior probabilities above 0.95 as likely to have experienced positive selection (Yang et al. 2005).

### Structural modeling

For genes whose protein structures have been resolved by X-ray crystallography or NMR, we downloaded the structures from the Protein Data Bank (PDB), prioritizing coverage of the structure to the translated nucleotide alignment. For genes whose protein structures were not resolved at the time of analysis, we then turned to predictive models from `AlphaFold` (Jumper et al. 2021). For structures not represented in either the PDB or `AlphaFold`, we performed homology modeling using templates with at least 40% sequence similarity using `Modeller` (Webb and Sali 2021). The resulting putative structures for each protein were compared by zDOPE, GA341 values, and RMSD values. Using these criteria in order, the best homology structure for each protein was selected and refined with `Modeller`’s DOPE-based loop-modeling protocols. For structures of proteins that did not contain regions that were represented in the nucleotide alignment, the ancestral representation of the alignment produced by `MASS-PRF` was used as a target for homology modeling using the original protein model as a template. The energy of the protein structure about the threaded region was then minimized with the `USCF Chimera` energy minimization protocol (Pettersen et al. 2004). `MASS-PRF` estimates of selection for each site within each gene were mapped to the corresponding protein models by translation and alignment of the `MASS-PRF` input sequence to the primary sequence of the homology model. The code for generating `MASS-PRF` structural heat maps is available at https://github.com/Townsend-Lab-Yale/massprf-protein-coloring.

## Results

A phylogeny of SARS-CoV-1, SARS-CoV-2, and SARS-like viruses isolated from non-human infections illustrates a history of infections of diverse mammalian hosts—hosts that in recent times have twice been the vector for zoonosis into humans (**Fig. 1A**). SARS-CoV-1 and SARS-CoV-2 constitute replicate zoonoses in distinct, moderately divergent coronavirus clades that enable replicated inference as to the selective histories associated with the coronavirus zoonotic process (**Fig. 1B**). The maximum-likelihood phylogenies produced by `RAxML` and `IQ-TREE 2` were equivalent in topology and nearly identical in branch lengths.

**Figure 1:**
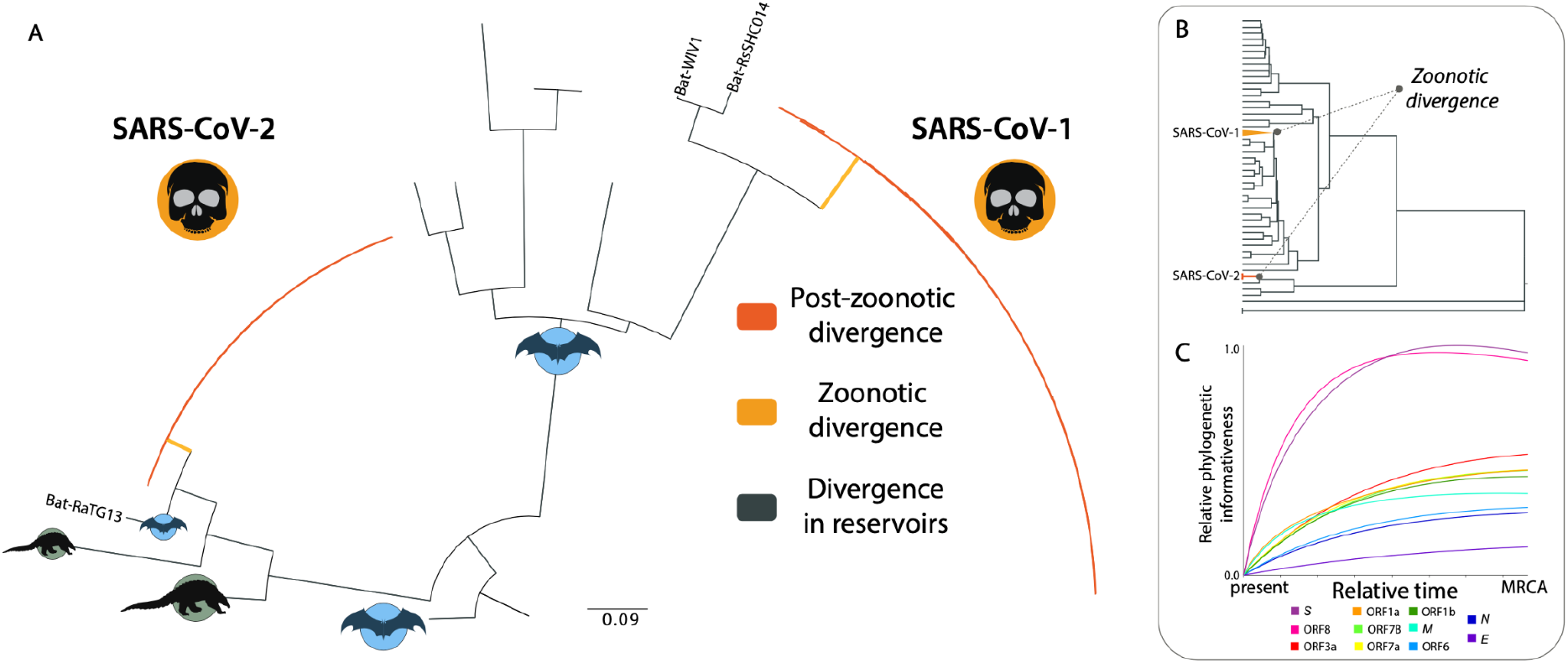
Molecular evolutionary tree, time tree, and phylogenetic informativeness profiles of 11 genes of SARS-CoV-1 and SARS-CoV-2. (**A**) Maximum-likelihood phylogeny of selected taxa from analysis of the *S*, ORF1b, ORF7a, *M*, ORF6, ORF3a, *E*, ORF8, *N*, ORF1a, and ORF7b genes of SARS-CoV-1 and SARS-CoV-2 and related SARS-like coronaviruses from nonhuman reservoir populations. (**B**) Time tree relative to the most recent common ancestor (MRCA) of the viruses studied. (**C**) Per-site phylogenetic informativeness of the *S*, ORF1b, ORF7a, *M*, ORF6, ORF3a, *E*, ORF8, *N*, ORF1a, and ORF7b genes of SARS-CoV-1 and SARS-CoV-2.

Genes exhibited divergent per-site phylogenetic informativeness (**Fig. 1C**), emblematic of gene- as well as site-specific rate distributions. The *S* and ORF8 genes exhibited rapid rises of phylogenetic informativeness (PI), a profile that is indicative of a substantial number of sites evolving at high rates of evolution. Lower phylogenetic informativeness across the divergences of interest was exhibited by ORF3a, ORF1a, ORF7B, ORF7a, ORF1b, *M*, ORF6, *N*, and lastly the very slowly-evolving *E* gene. Regardless of the slope of increase of informativeness, all genes peak in informativeness at or near the base of the inferred phylogeny, indicating minimal potential for phylogenetic noise (homoplasy) that might compromise phylogenetic inference (Townsend 2007; Townsend and Leuenberger 2011; Dornburg et al. 2019; **Fig. 1B–C**).

Evaluations of selection after zoonosis and during the early spread of SARS-CoV-2 using `PAML` revealed additional sites in the *S* gene of SARS-CoV-2 that were under selection, along with some sites within *M* and ORF3a (**Supplementary Fig. 1; Supplementary Table S1**). In contrast, numerous sites of the *S* gene and other genes were under positive selection in SARS-CoV-1. Application of `MASS-PRF` to evaluate selective pressures proximate to zoonoses revealed abundant purifying selection and evolution under neutrality (**Supplementary Fig. 1**). Numerous genes (60%) with low levels of sequence divergence and polymorphism could not be analyzed with the PRF model as a consequence of their low levels of sequence divergence and polymorphism (**Supplementary Table S1**). Their paucity of both variation and divergence is consistent with strong purifying selection (Zhao et al. 2017).

Estimates of the scaled selection coefficient 𝛄 during the zoonotic branch rose above zero for a few regions of the S proteins in SARS-CoV-2 and SARS-CoV-1 (**Fig. 2**). Several amino acids in the short N-Terminal Domain of SARS-CoV-2 were found to exhibit a signature of positive selection post-zoonosis (**Fig. 2A**). Several amino acids in this same region of the SARS-CoV-1 S protein also exhibited statistically significantly positive 𝛄 estimates that correspond to a region within the globular head (**Fig. 2B**). In contrast, the vast majority of regions in the *S* gene of both viruses exhibited 𝛄 values statistically significantly below zero during the zoonotic branch, indicative of strong purifying selection. This purifying selection was substantially indicated in both viruses, but with even greater power (tighter confidence) across the SARS-CoV-2 S protein than across the SARS-CoV-1 S protein. The S2 subunit of SARS-CoV-1 exhibited notably higher model uncertainty than the S1 subunit (**Fig. 2B**). The only exception to this general pattern of higher uncertainty for SARS-CoV-1 relative to SARS-CoV-2 was one region of the primary protein structure in S2 near the S cleavage site. This region exhibited a striking pattern of synonymous and replacement polymorphism and divergence in SARS-CoV-2, with high model uncertainty overlapping 𝛄 = 0 such that it does not constitute a statistically significantly positive 𝛄 estimate (**Fig. 2A**).

**Figure 2.**
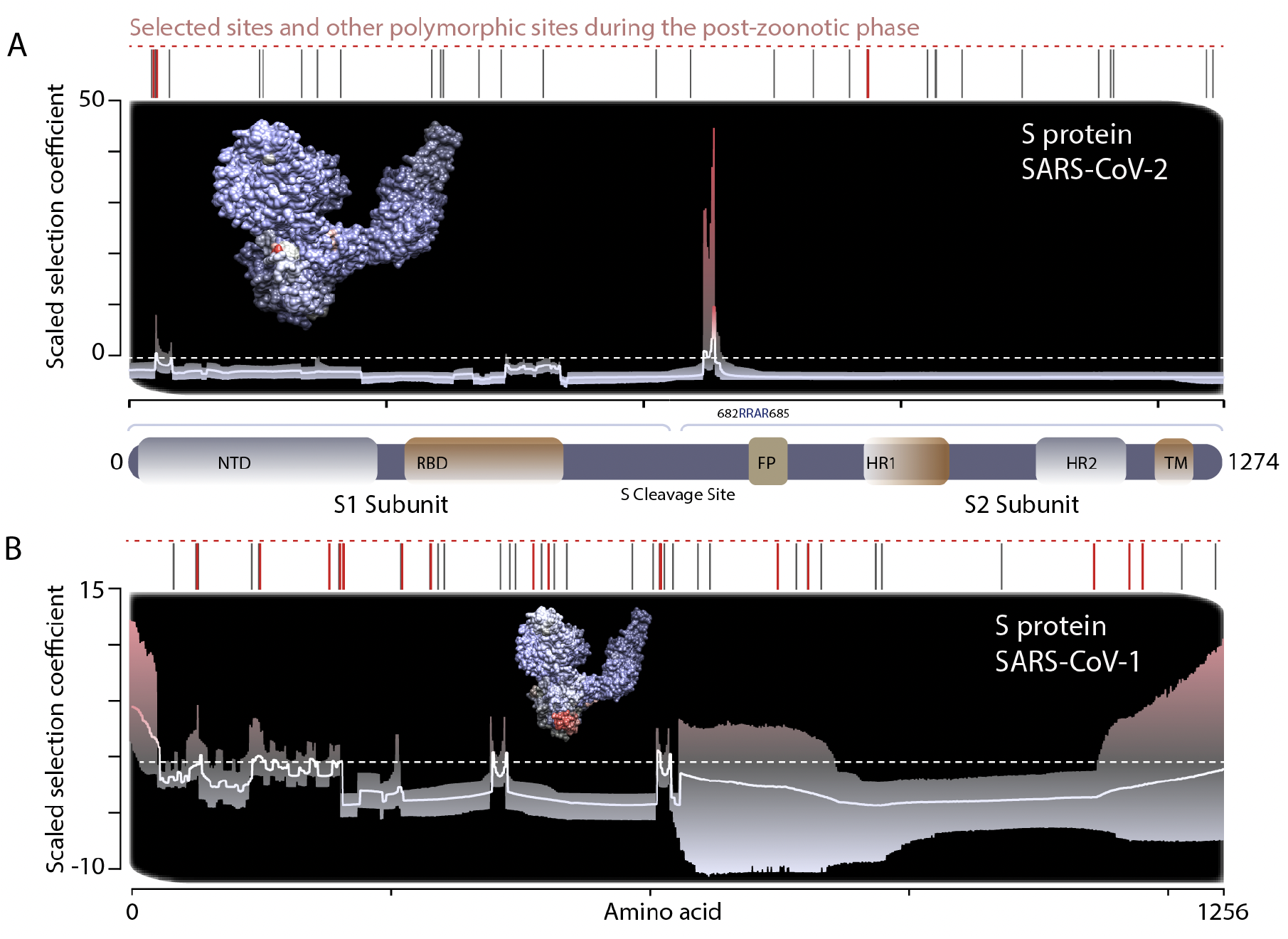
The regional strength of selection for amino-acid changes along the zoonotic branch of the *S* protein of (A) SARS-CoV-2 and (B) SARS-CoV-1. Model-averaged regional scaled selection across the primary protein structure is indicated by the white line (compared to a scaled selection coefficient of zero, dashed white line; coefficients greater than zero indicate positive, adaptive selection and coefficients less than zero indicate negative, purifying selection; 95% model interval: light gray to red gradient). (*inset*) S-protein tertiary structure models with amino acids colored along a blue→red color gradient, scaled to reflect the negative→positive selection gradient. Individual polymorphic sites (gray hashes) and sites identified by PAML as being under selection during viral spread in humans subsequent to zoonosis (red hashes) are indicated above each plot. The N-Terminal Domain (NTD, gray), Receptor-Binding Domain (RBD, brown), Fusion Peptide (FP), Heptad-Repeat domain 1 (HR1), Heptad-Repeat domain 2 (HR2), and TransMembrane (TM) domains are diagrammed in relation to the S-protein primary structure.

Substantial signal of extensive purifying selection proximate to zoonosis similar to those observed in the S protein of SARS-CoV-1 and SARS-CoV-2 manifests in all proteins expressed by both coronavirus genomes (**Supplementary Figures 2–3**; **Supplementary Table S1**). For example, the non-structural protein NSP3 in the SARS-CoV-2 ORF1a polyprotein-encoding gene exhibited estimates of the scaled selection coefficient that were significantly less than zero (**Fig. 3A**). A similar signature of pervasive purifying selection proximate to zoonosis was obtained for the N protein (**Fig. 3B**) and ORF3a protein of SARS-CoV-2 (**Fig. 3C**). Similarly, we find evidence for purifying selection in the non-structural proteins NSP12 of ORF1b and NSP 5 of the ORF1a polyprotein-encoding gene (**Fig. 3D–E**) in SARS-CoV-1. Other results for other genes demonstrate similar patterns of strong purifying selection across the genome (**Supplementary Figs. 2–3**; **Supplementary Table S1**). Collectively, these results reveal that proximate to the time of zoonosis, molecular evolution in the genomes of both viruses was dominated by purifying selection and neutral evolution.

**Figure 3.**
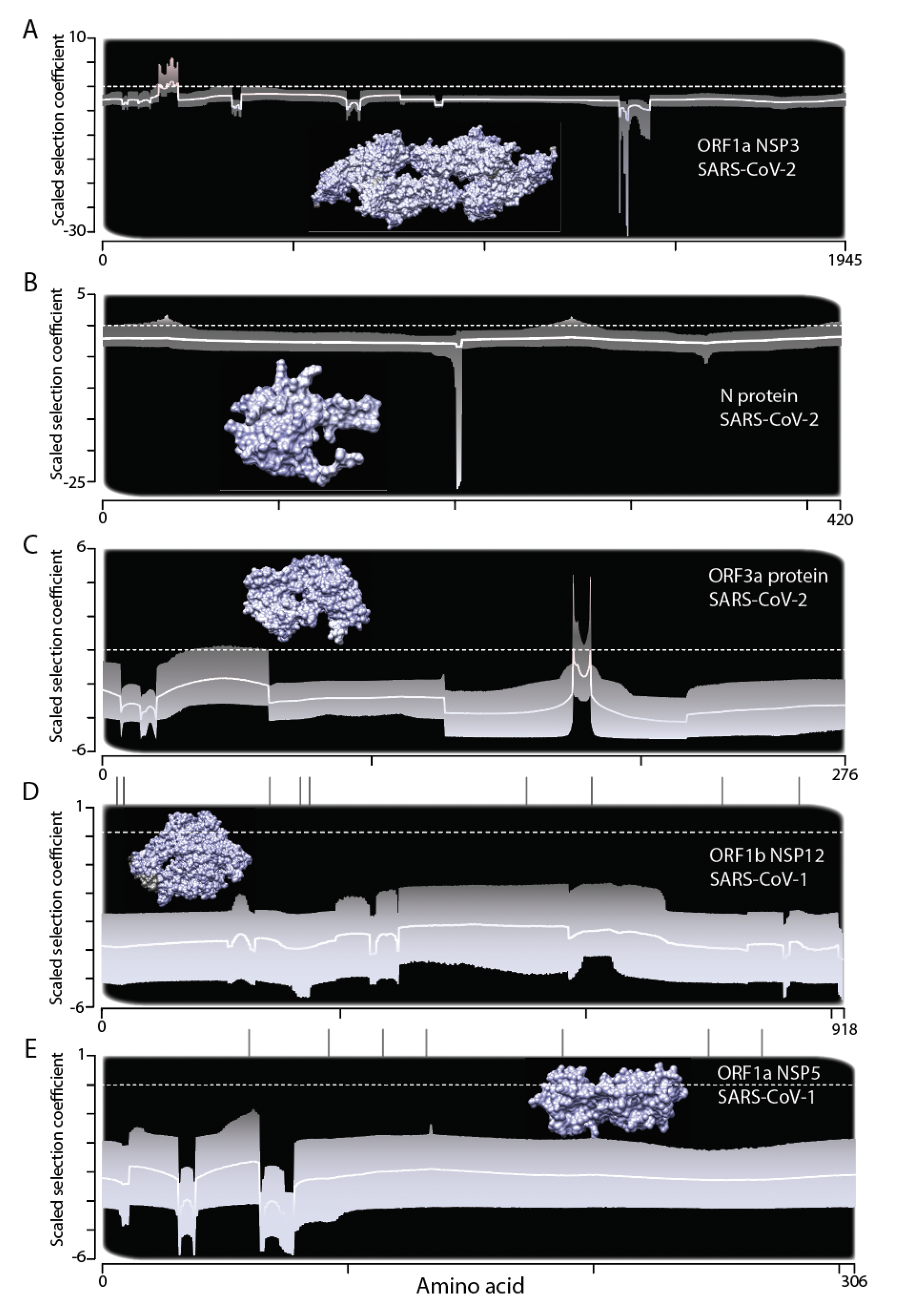
Evidence for purifying selection proximate to zoonosis across (A) ORF1a NSP3, (B) N, and (C) ORF3a in SARS-CoV-2, and (D) ORF1b NSP12 and (E) ORF1a NSP5 in SARS-CoV-1. Model-averaged regional scaled selection across the primary protein structure is indicated by the white line (compared to a scaled selection coefficient of zero, dashed white line; coefficients greater than zero indicate positive, adaptive selection and coefficients less than zero indicate negative, purifying selection; 95% model interval: light gray to dark gray gradient). (*inset*) Models of the tertiary structure of each protein, with amino acids colored along a blue→red color gradient, scaled to reflect the negative→positive selection gradient. Individual polymorphic sites (gray hashes) are indicated above each plot.

In contrast to the strong signature of purifying selection proximate to zoonosis across most of the SARS-CoV-1 and SARS-CoV-2 genomes (e.g. **Fig. 3**), we find evidence for positive selection in Orf7a for both SARS-CoV-2 and SARS-CoV-1 (**Fig. 4A–B**). In SARS-CoV-2, our analysis of the ORF7a protein reveals strong statistical support for positive selection proximate to zoonosis in the distinctive immunoglobulin (Ig)-like ectodomain (positions 71 and 73; Zhou et al. 2021) and in the hydrophobic transmembrane domain (positions 110, 111, and 115; **Fig. 4A**). Some variants in this transmembrane domain have been shown to confer increased protein stability through allosteric effects (Lobiuc et al. 2021). A distinct region of the ectodomain in SARS-CoV-1is under positive selection along the zoonotic branch (positions 23–25; **Fig. 4B**). The protein Orf1b NSP12 has been referred to as the most highly conserved protein in coronaviruses (Subissi et al. 2014);nevertheless; it exhibited evidence of selection in SARS-CoV-2 (**Fig. 4C**), but not in SARS-CoV-1 (**Fig. 3D**).

**Figure 4.**
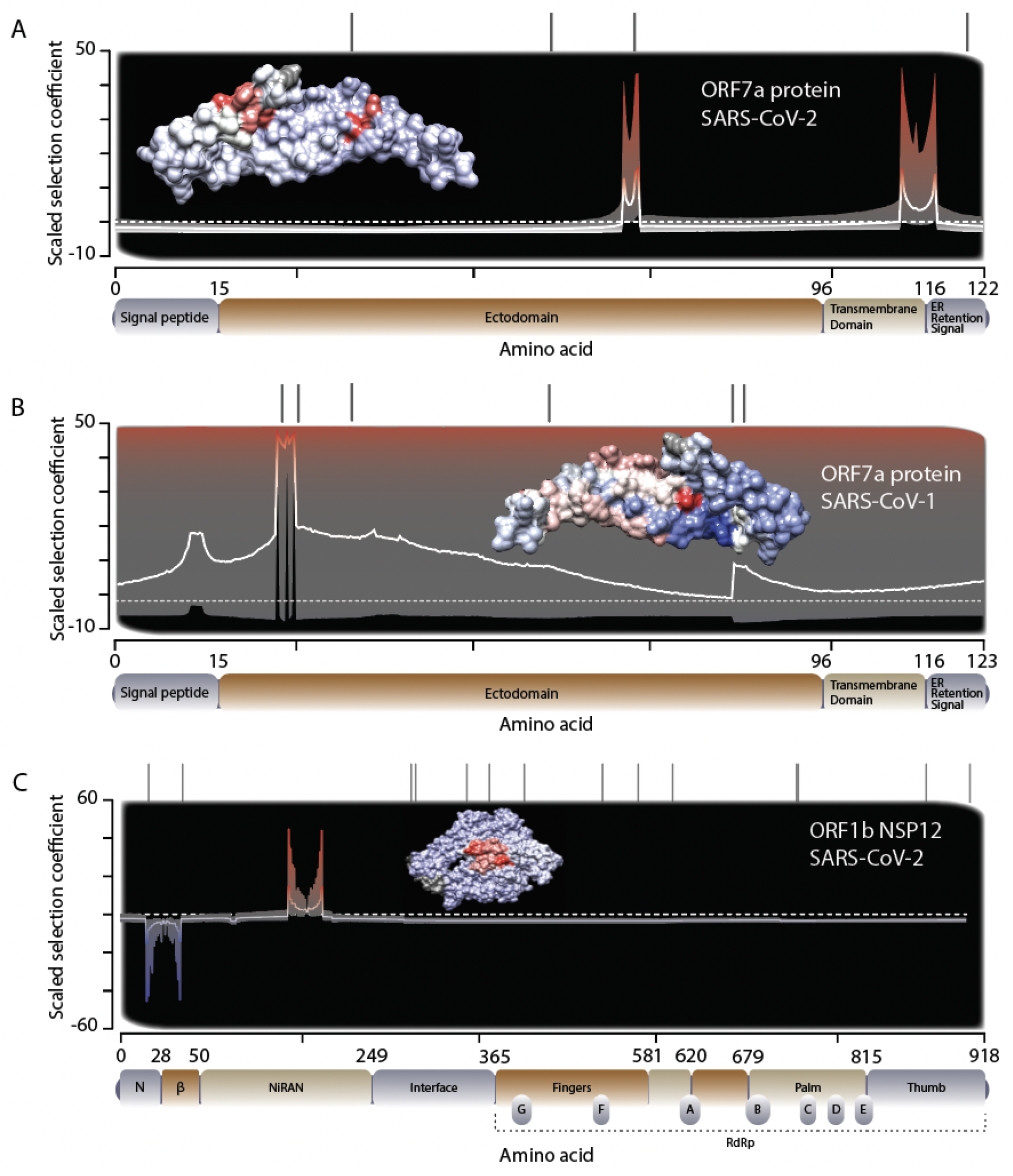
Evidence for selection in SARS-CoV-2 proximate to zoonosis within (A) ORF7a of SARS-CoV-2, (B) ORF7a of SARS-CoV-1, and (C) NSP12 within ORF1b in SARS-CoV-2. Model-averaged regional scaled selection across the primary protein structure is indicated by the white line (compared to a scaled selection coefficient of zero, dashed white line; coefficients greater than zero indicate positive, adaptive selection and coefficients less than zero indicate negative, purifying selection; 95% model interval: the light-gray to reddish gradient). (*insets*) Models of the tertiary structure of each protein with amino acids colored along a blue→red color gradient scaled to reflect the corresponding negative→positive selection gradient. Individual polymorphic sites (gray hashes) are indicated above each plot.

## Discussion

Here we have illuminated the molecular evolutionary history spanning the zoonosis of SARS-CoV-2 and SARS-CoV-1. These viruses have independently been transmitted from natural reservoirs to humans within the last two decades, providing replicated and highly informative phylogenetic datasets for inferences of selection. The molecular evolutionary rates of genes within SARS-like coronaviruses can be grouped in three clusters, with the Spike gene (*S*) and ORF8 exhibiting the largest numbers of faster-evolving sites, numerous other genes with sites evolving at a range of moderate rates, and *E* exhibiting an exceptionally low rate of molecular evolution. Proximate to zoonosis, evolution of proteins in SARS-CoV-1 and SARS-CoV-2 was characterized almost exclusively by purifying selection and neutral evolution, with positive selection detectable in only a few sites in the genome. This finding proximate to zoonosis contrasts sharply with the ongoing spread of SARS-CoV-2 that is characterized by the appearance of new genetic variants, many of which confer immuno-evasive or other properties that are subject to positive selection (D.P. Martin et al. 2021; Harari et al. 2022; Martin et al. 2022; C.W. Tan et al. 2022). Collectively, these results indicate that zoonosis is likely mediated by chance interactions with wildlife that is not contingent on rapid viral evolution for transmission to humans (MacLean et al. 2021; C.C.S. Tan et al. 2022). These results underscore the importance to global public health of careful surveillance for emerging disease threats and the importance of rapid and effective responses.

Most attention regarding molecular adaptation in SARS viruses has been devoted to the antigenically variable regions of *S* (Zhang et al. 2006; Chan and Zhan 2021; Kang et al. 2021; MacLean et al. 2021; Salleh et al. 2021). Indeed, insertions in the zoonotic lineage have been reported to have functional importance (Gussow et al. 2020; Chan and Zhan 2021). Our results reveal moderate, asymmetric selection patterns between the S protein in SARS-CoV-1 and SARS-CoV-2 proximate to zoonosis. Our analysis identified a small region within the S protein of SARS-CoV-1 that exhibited a signal of positive selection during zoonosis that exhibited no substantial or significant selection in SARS-CoV-2, a finding that parallels widespread selection on sites in the S protein of SARS-CoV-1, and to a lesser extent, SARS-CoV-2, subsequent to zoonosis. These findings are consistent with other studies of selection that have identified immuno-evasive mutations during the pandemic spread of SARS-CoV-2 (Lo Presti et al. 2020; Rochman et al. 2021; Kistler et al. 2022) as well as studies of selection in the most prevalent human coronavirus, HCoV-OC43 (Ren et al. 2015).

The Pfizer and ModeRNA vaccines target the Spike protein of SARS-CoV-2, which presents a surface that antibodies will easily encounter. It was a propitious target for the first vaccines, which had to be developed and deployed quickly; the extracellular portions of the membrane (M) and envelope (E) surface proteins are somewhat shielded by the Spike (Bianchi et al. 2020). However, our analyses complement a host of immunological studies demonstrating that mutations in the *S* gene can encode amino-acid changes that confer substantially divergent epitope structure with each mutation. Indeed, over the course of the COVID-19 pandemic, individual amino-acid changes in the Spike protein conferred selective advantages that enabled immune evasion (Korber et al. 2020; Weisblum et al. 2020; Greaney et al. 2021). Therefore, future generations of vaccine development could include as targets the membrane (M) and envelope (E) proteins, which our results reveal to be slower evolving (**Fig. 1**) due to substantial purifying selection (**Supplementary Figure 1**). A high Ig response against both S and M proteins has been observed in patients with severe COVID-19 (S. Martin et al. 2021). A vaccine that elicits an antigenic response to a conserved portion of S (Campbell et al. 2023), M, or E might provide much more general and durable protection, instead of the short duration of protection we can count on from a generalized Spike-targeting vaccine (Townsend et al. 2021; Townsend et al. 2023).

The zoonotic branch of the viral phylogeny where we have analyzed the signal of selection is likely to predominantly reflect viral evolution on non-human hosts, followed by only a limited number generations of transmission from human to human, given the rapidity with which SARS-CoV-2 was identified. However, selection in those first human hosts may have been strong. Rather than finding that the zoonosis of SARS-CoV-2 was dominated by changes in the *S* gene, we found a strong signal of selection in NSP12. Accordingly, a substitution in NSP12 has recently been highlighted as providing a selective advantage to transmission during the evolution of SARS-CoV-2 variants (Goldswain et al. 2023). Our results indicate additional sites that were selected proximate to zoonosis that might have been important for initial human transmissibility.

An additional site of adaptation occurred along the zoonotic branches in ORF7a, which may have played an underappreciated role in the zoonosis of both SARS-CoV-1 and SARS-CoV-2. *In vitro* experiments have shown that expression of ORF7a protein from SARS-CoV-1 is associated with the enhanced production of pro-inflammatory cytokines (Kanzawa et al. 2006); ORF7a of SARS-CoV-2 has been attributed a role in inhibition of interferon signaling and limitation of the activation of mechanisms that restrict viral replication (Xia et al. 2020; Cao et al. 2021). ORF7a has also been demonstrated to bind to CD14+ monocytes and has been proposed to limit the ability of these monocytes to express antigens (Zhou et al. 2021). More recently, ORF7a has been shown to block SERINC5, which otherwise inhibits virus-cell fusion (Timilsina et al. 2022), and down-regulates MHC class-I surface expression (Arshad et al. 2022; Zheng et al. 2022). The adaptive substitutions we have noted in ORF7a were fixed within each SARS virus proximal to zoonosis, either while propagating in reservoir species or during the period spanning the first human transmissions. Given that evidence strongly suggests bats are the primary reservoirs of SARS-like coronaviruses (Li et al. 2005), the bat immune system is a likely source of pre- and proximal-zoonotic selective pressure. Notably, bat species have been shown to have constitutive expression of type-I interferon (Zhou et al. 2016), which may select for viral species that can strongly manipulate interferon signaling, leading to high pathogenicity when introduced to humans (Schountz et al. 2017).

Purifying selection was abundant across the genomes of SARS-CoV-1 and SARS-CoV-2 proximate to zoonosis. However, there is also evidence for selection in gene regions associated with host immune evasion and viral replication in both viral lineages. Awareness of these genetic adaptations proximate to zoonosis is a key component of efforts to determine general rules underlying the ability of coronaviruses to switch hosts. Further molecular evolutionary study of additional zoonotic events, is warranted—whether they lead to a pandemic or not—to provide continued illumination of adaptations that enable zoonosis. Understanding these rules may help in preventing the next pandemic.

## Acknowledgements

We thank undergraduate Jaiveer Singh for contributions to our initial forays into research on this topic, and to Naser Hatami for some edits incorporated into the final draft. We are grateful to the Yale Center for Research Computing for the use of the research computing infrastructure. Vector art in the supplemental materials was modified from free-to-use vector graphics available from vecteezy.com (human silhouettes:

https://www.vecteezy.com/free-vector/free-family-silhouette and all-free-download.com; bat silhouette:

https://all-free-download.com/free-vector/download/halloween-design-elements-bats-icons-silhouettes-design_288876.html)

## Data availability

Data and code are publicly available. Input and output files for MASS-PRF, PAML, phylogenies, protein structure, and phylogenetic informativeness are available on Zenodo at DOI:10.5281/zenodo.7953678.

