## Supplemental materials for "Purifying selection and adaptive evolution proximate to the zoonosis of SARS-CoV-1 and SARS-CoV-2"

**Contents:**

**Table S1:** Positively selected sites among SARS coronaviruses (MASS-PRF zoonosis and PAML post-zoonosis)

**Figure S1**: Putative zoonotic and post-zoonotic selection on sites of 11 genes in SARS-CoV-1 and SARS-CoV-2.

**Figure S2:** Evidence for purifying selection proximate to zoonosis in SARS-CoV-2.

**Figure S3**: Evidence for purifying selection proximate to zoonosis in SARS-CoV-1.

**Table S1**. Positively selected sites among SARS coronaviruses (MASS-PRF zoonosis and PAML post-zoonosis)

| | MASS-PRF | | PAML | |
|---|---|---|---|---|
| Gene | SARS1 | SARS2 | **SARS-CoV-1** | **SARS-CoV-2** |
| *E* | [a] | [b] | **14 (0.929)** | None |
| *S* | **1–12** | None | **75 (0.971*), 147 (0.998**), 227 (0.969*), 239 (0.971*), 243 (0.969*), 244 (0.971*), 311 (0.972*), 344 (0.970*), 462 (0.971*), 479 (0.997**), 608 (0.971*), 609 (0.972*), 743 (0.972*), 778 (0.970*), 1107 (0.969*), 1148 (0.971*), 1163 (0.970*)** | **28 (0.896), 32 (0.891), 860 (0.967*), 861 (0.899)** |
| *N*[c] | [d] | None | None | None |
| *M* | [f] | [d] | **11 (0.943)** | **1 (0.804)** |
| Orf7b | [a,d,e] | [a] | None | None |
| Orf7a | **23, 24, 25** | **71, 72, 73, 110, 111, 115** | None | None |
| Orf8 | [g] | [a,b,d] | **17 (0.943), 31 (0.943), 48 (0.940)** | None |

| | | | | |
|---|---|---|---|---|
| Orf6 | [a,b,e] | [a,d,e] | None | None |
| Orf3a | [b] | None | **12 (0.986*), 26 (0.896), 82 (0.985*), 143 (0.886)** | **270 (0.816)** |
| nsp1 | [a] | [e] | **33(0.984*), 82(0.984*), 95(0.984*), 130(0.984*)** | None |
| nsp2 | None | **95** | **291(0.846)** | None |
| nsp3 | None | None | **14(0.925), 845(0.923), 1082(0.924), 1404(0.923)** | None |
| nsp4 | None | None | **6(0.966*), 204(0.966*), 231(0.966*), 307(0.966*)** | None |
| nsp5 | None | [d,e] | None | None |
| nsp6 | [d,e] | [d] | None | None |
| nsp7 | [a,d] | [d,e] | **7(0.962*), 68(0.962*)** | None |
| nsp8 | [d] | [d] | None | None |
| nsp9 | [a,d] | [d,e] | **37(0.977*), 67(0.977*), 76(0.977*)** | Insufficient polymorphism |
| nsp10 | [d,e] | [d] | None | None |
| nsp11 | [a,b,d,e] | [a,b,d,e] | None | None |
| nsp12 | None | **61,73** | **28(0.850)** | None |

| | | | | |
|---|---|---|---|---|
| nsp13 | [d] | [d] | None | None |
| nsp14 | None | None | None | None |
| nsp15 | None | None | None | None |
| nsp16 | None | [d] | None | None |

[a] insufficient power—synonymous polymorphism (SP) was too low,
[b] insufficient power—synonymous divergence (SD) was too low,
[c] two sequences were removed from the *N* gene of SARS-CoV-1 due to exhibiting a frameshift mutation therein.
[d] insufficient power—replacement divergence (RD) was too low,
[e] insufficient power—replacement polymorphism (RP) was too low.
[f] Running MASS-PRF on the SARS-CoV-1 *M* gene consistently failed due to the cluster modeling for this gene being too diffuse, requiring exponentially too much memory
[g] ORF8 of SARS-CoV-1 frequently featured deletions that disrupted the coding sequence. We were therefore not able to run MASSPRF on this polymorphism data.

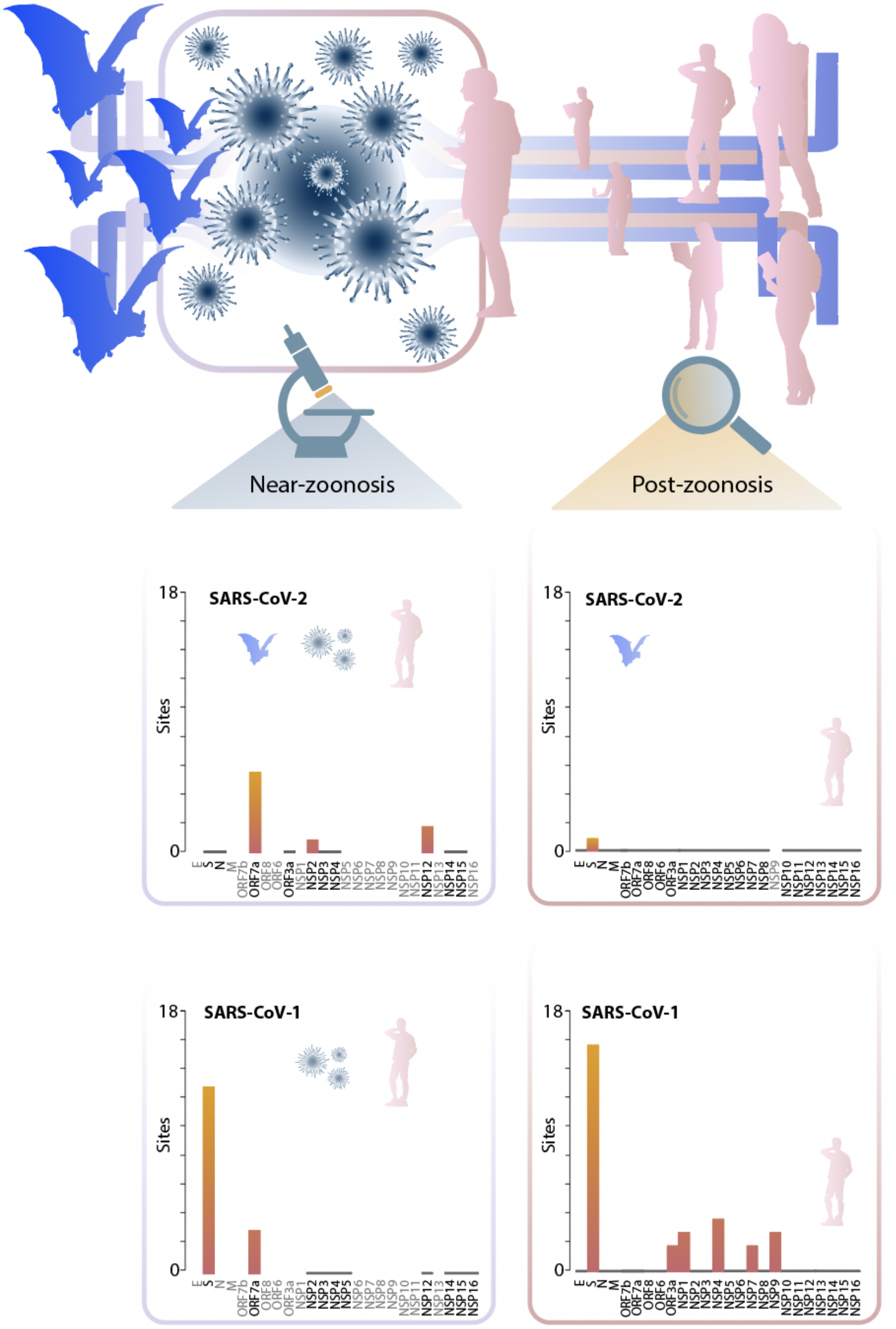


**Figure S1**. Putative zoonotic and post-zoonotic selection on sites of 11 genes in SARS-CoV-1 and SARS-CoV-2. Genes and NSPs with levels of sequence divergence too low for analysis are indicated with gray text.

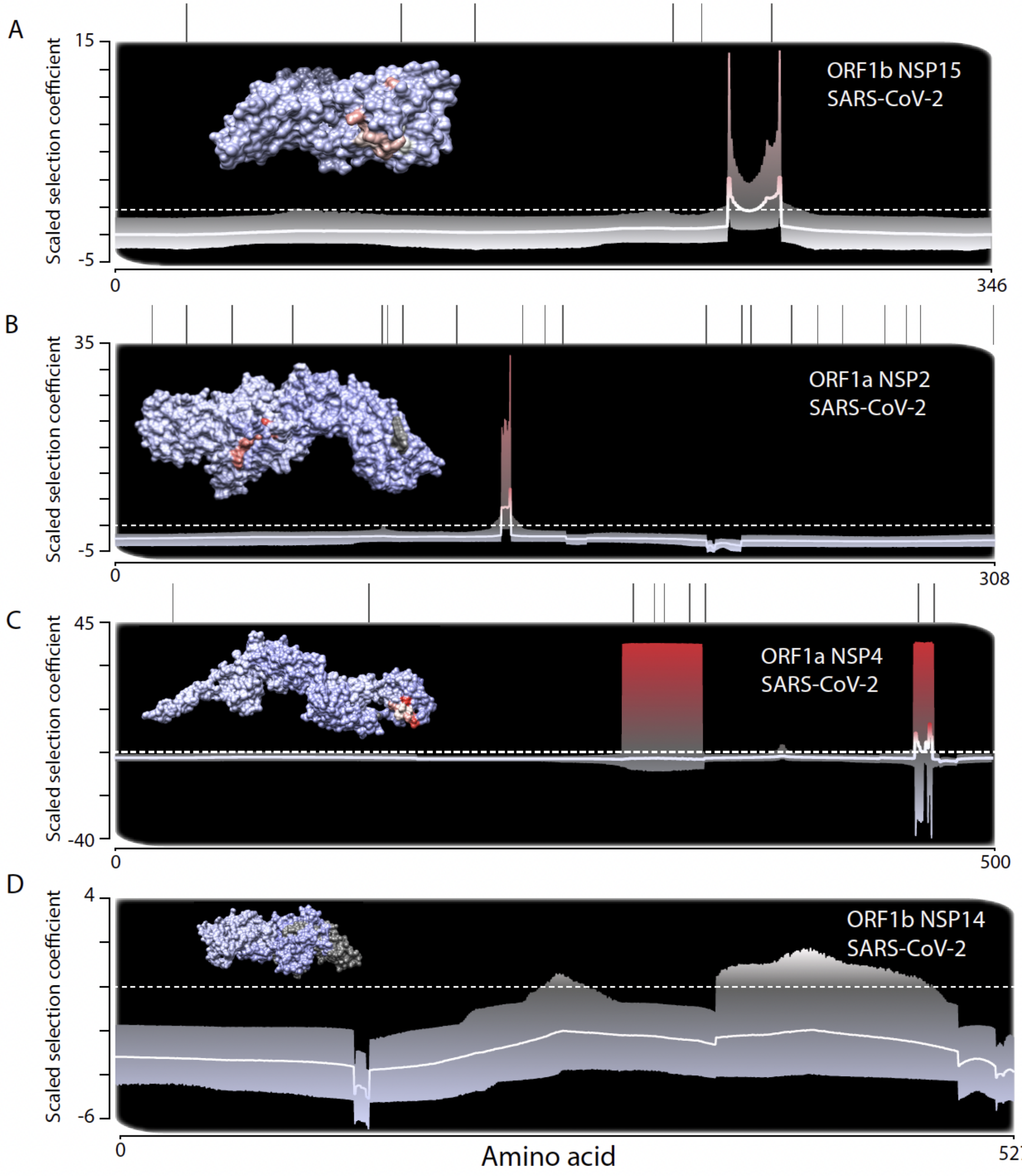


**Figure S2**. Evidence for purifying selection proximate to zoonosis across (**A**) ORF1b NSP15, (**B**) ORF1a NSP2, (**C**) ORF1a NSP4, and (**D**) ORF1b NSP14 in SARS-CoV-2. Model-averaged regional selection across the primary protein structure is indicated by the white line (compared to a scaled selection coefficient of zero, dashed white line; coefficients greater than zero indicate positive, adaptive selection and coefficients less than zero indicate negative, purifying selection; 95% model interval: light→dark gray gradient). (*inset*) Models of the tertiary protein structure of each gene (blue→red color: negative→positive selection). Polymorphic sites (gray hashes) and sites identified by PAML as being under selection during viral spread in humans subsequent to zoonosis (red hashes) are indicated above each plot.

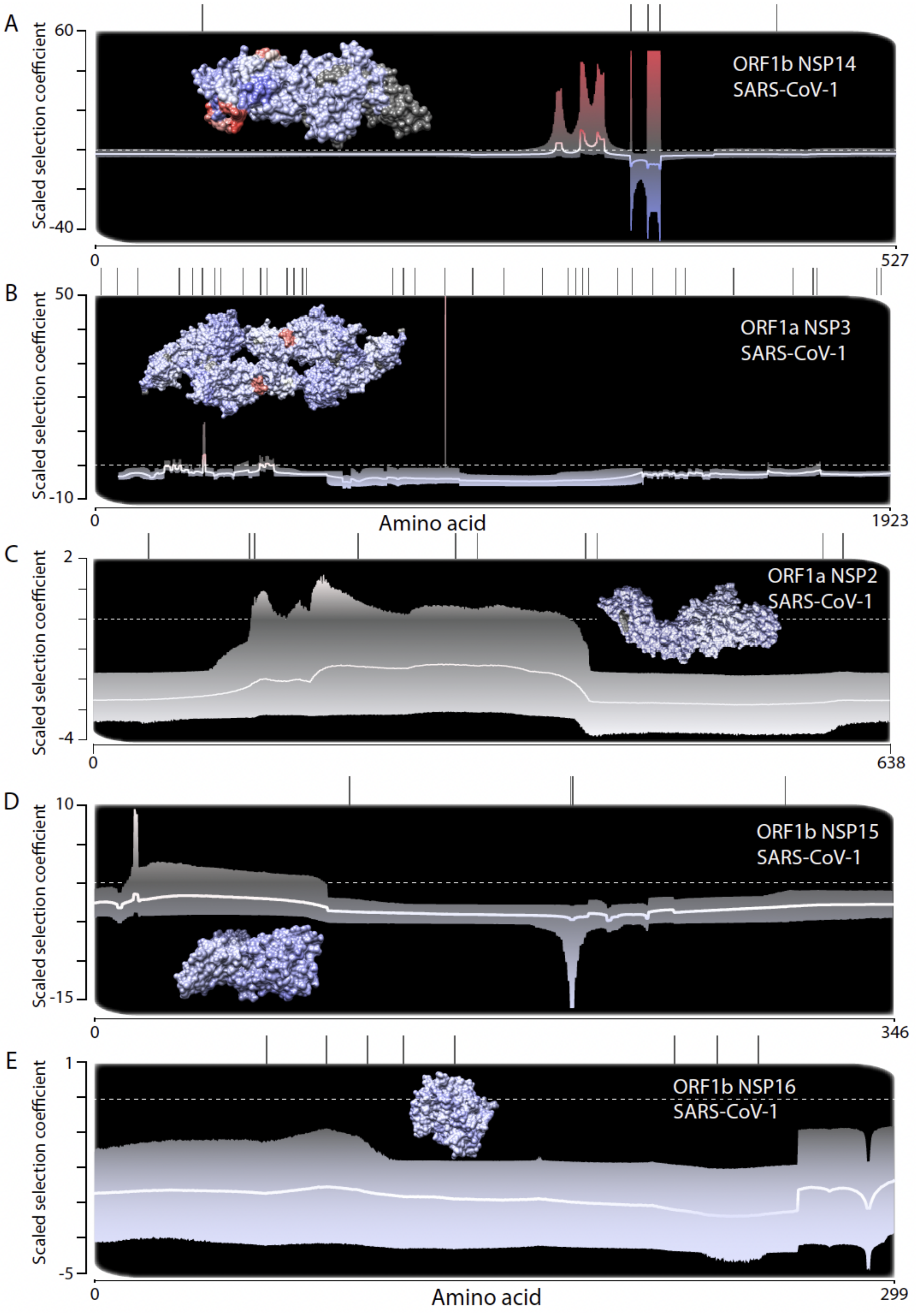
A
Scaled selection coefficient
60
-40
0
527
ORF1b NSP14
SARS-CoV-1
B
50
-10
1923
ORF1a NSP3
Amino acid
C
2
-4
638
ORF1a NSP2
D
10
-15
346
ORF1b NSP15
E
1
-5
299
ORF1b NSP16

**Figure S3.** Evidence for purifying selection proximate to zoonosis across (**A**) ORF1b NSP14, (**B**) ORF1a NSP3, (**C**) ORF1b NSP2, (**D**) ORF1b NSP15, and (**E**) ORF1b NSP16 in SARS-CoV-1. Model-averaged regional selection across the primary protein structure is indicated by the white line (compared to a scaled selection coefficient of zero, dashed white line; coefficients greater than zero indicate positive, adaptive selection and coefficients less than zero indicate negative, purifying selection; 95% model interval: light gray to dark gray gradient). (*inset*) Models of the tertiary protein structure of each gene (blue→red color: negative→positive selection). Individual polymorphic sites (gray hashes) and sites identified by PAML as being under selection during viral spread in humans subsequent to zoonosis (red hashes) are indicated above each plot.
